# Night watch during REM sleep for the first-night effect

**DOI:** 10.1101/668020

**Authors:** Masako Tamaki, Yuka Sasaki

**Author notes:** CORRESPONDING AUTHOR: Yuka Sasaki.

## Abstract

We experience disturbed sleep in a new place, and this effect is known as the first-night effect (FNE) in sleep research. We previously demonstrated that the FNE was associated with a protective night-watch system during NREM sleep in one hemisphere, which is shown as interhemispheric asymmetry in sleep depth in the default-mode network (DMN), and interhemispheric asymmetry in increased vigilance to monitor external stimuli. The present study investigated whether rapid eye movement (REM) sleep exhibited a form similar to a night-watch system during NREM sleep. First, we tested whether theta activity, which is an index of the depth of REM sleep, showed interhemispheric asymmetry in association with the FNE, by source-localizing to the DMN. However, interhemispheric asymmetry in theta activity during REM sleep was not found in association with the FNE. Next, we tested whether vigilance, as measured by evoked brain responses to deviant sounds, was increased in one hemisphere and showed interhemispheric asymmetry in association with the FNE during REM sleep. Because vigilance is different between the phasic period where rapid eye movements occur and the tonic period where rapid eye movements do not occur during REM sleep, REM sleep was split into phasic and tonic periods for measurements of evoked brain responses. While the evoked brain responses are generally small during the phasic period without the FNE, we found that the evoked brain response was significantly augmented by the FNE during the phasic period. In contrast, the evoked brain response during the tonic period did not differ by the presence of the FNE. Interhemispheric asymmetry in brain responses was not found during the phasic or tonic periods. These results suggest that a night-watch system for the FNE appears as interhemispheric asymmetry in sleep depth and vigilance during NREM sleep, but it appears as increased vigilance in both hemispheres during the phasic period, when vigilance to external stimuli is generally reduced without the FNE, during REM sleep. Therefore, a night-watch system associated with the FNE may be subserved by different neural mechanisms during NREM and REM sleep.

## Introduction

Sleep is crucial for the maintenance of daily life (Stickgold, 2005; Imeri and Opp, 2009). The psychological and behavioral consequences of declines in sleep quality may be severe (Carskadon and Dement, 1981; Drummond et al., 2000; Buysse et al., 2011; Okawa, 2011; Czeisler, 2013). However, sleep may be decreased as a protective mechanism under specific circumstances (Rattenborg et al., 1999; Peever and Fuller, 2017). One form of this mechanism is the first-night effect (FNE), which is widely known in human sleep research (Agnew et al., 1966; Carskadon and Dement, 1979; Tamaki et al., 2005a; Tamaki et al., 2005b; Tamaki et al., 2014; 2016). The FNE is a temporary sleep disturbance that occurs specifically in the first session of sleep experiments in young healthy adults, and it manifests as prolonged sleep-onset latency, frequent arousals, and decreased deep non-rapid eye movement (NREM) sleep (Roth et al., 2005). Our previous study demonstrated that the FNE was not merely a sleep disturbance but a manifestation of a protective night-watch system in one hemisphere, which is less asleep and more vigilant than the other hemisphere to monitor unfamiliar surroundings during deep NREM sleep (Tamaki et al., 2016). More specifically, slow-wave activity, which is an index of sleep depth during NREM sleep, decreases in the left hemisphere compared to the right hemisphere regionally in the default-mode network (DMN) on day 1 with the FNE, which causes interhemispheric asymmetry in slow-wave activity. The amplitude of an evoked brain potential during deep NREM sleep correlates with vigilance, and it was increased in one hemisphere on day 1, which caused interhemispheric asymmetry in vigilance. These interhemispheric asymmetries in local sleep depth and vigilance to monitor the external world function as a night-watch system, which may counteract vulnerability during deep NREM sleep (Tamaki et al., 2016).

In contrast to NREM sleep, whether the FNE influences brain activities during rapid eye movement (REM) sleep, and its possible mechanisms, are less well understood. Approximately 30 studies investigated the FNE during REM sleep in healthy young adults. The characteristics of the FNE during REM sleep include decreased time spent in REM sleep (Rechtschaffen and Verdone, 1964; Kales et al., 1967b; Sforza et al., 2008), delayed onset of REM sleep (Agnew et al., 1966; Kales et al., 1967b), and increased micro-arousals during REM sleep (Sforza et al., 2008). To our best knowledge, only two papers (Toussaint et al., 1997; Curcio et al., 2004) investigated brain activities quantitatively during REM sleep. However, the results from these studies are contradictory (Toussaint et al., 1997; Curcio et al., 2004). One study showed that the FNE decreased electroencephalography (EEG) power in wide ranges of frequency bands, including delta (0.5-3.5 Hz), theta (4-7.5 Hz), and beta (13-21.5 Hz) bands (Toussaint et al., 1997). The other study reported that the FNE increased theta power (5 Hz) (Curcio et al., 2004). No study investigated brain activities separately for each hemisphere or whether the FNE altered vigilance during REM sleep. Therefore, whether the FNE alters sleep depth or vigilance level during REM sleep to act as a night-watch system in an analogous manner to NREM sleep is not known.

Notably, it is suggested that the ability to process external stimuli differs depending on whether rapid eye movements appear (phasic period) or not (tonic period) during REM sleep (Price and Kremen, 1980; Sallinen et al., 1996; Takahara et al., 2002; 2006b; Wehrle et al., 2007). First, the amplitudes of evoked brain responses differ significantly between the phasic and tonic periods. That is, much smaller brain responses are observed during the phasic than the tonic period (Sallinen et al., 1996; Takahara et al., 2002; 2006b). The amplitudes of evoked potentials correlate with the degree of vigilance (Nielsen-Bohlman et al., 1991; Michida et al., 2005). Therefore, the finding of smaller potentials during the phasic period suggests that the degree of vigilance to external stimuli is significantly decreased when the eyes are moving during the phasic compared to tonic periods during REM sleep without the FNE. Second, one study investigated blood oxygenation level-dependent (BOLD) responses to acoustic stimulations (Wehrle et al., 2007) and reported that acoustic stimulations during the tonic period induced BOLD activations in the auditory cortex to some degree. In contrast, BOLD activations in the auditory cortex during the phasic period were only minimal. Therefore, the ability to process external stimuli may be reduced significantly during the phasic period but sustained to some extent during the tonic period. Therefore, it is important to split REM sleep into phasic and tonic periods for examination of evoked brain responses during REM sleep.

The present study investigated whether the FNE affected the sleep depth and vigilance of REM sleep, and whether decreased sleep depth or enhanced vigilance, if any, shows interhemispheric asymmetry in an analogous manner to NREM sleep. Theta activity was investigated as an index of the depth of REM sleep because it is one of the major spontaneous brain oscillations during REM sleep (Takahara et al., 2006a). Our previous study suggested that brain activities may need to be source-localized using individual anatomical brain information for the detection of asymmetry in sleep depth (Tamaki et al., 2016). Moreover, interhemispheric asymmetry in sleep depth is difficult to identify on the sensor space, at least in the case of NREM sleep (Tamaki et al., 2016). Therefore, we tested whether the FNE during REM sleep involved interhemispheric asymmetry in theta activity in the DMN using a source-localization technique that combines EEG, structural MRI, and PSG in the sleeping brain. To test whether the FNE altered vigilance, we measured evoked brain responses to external stimuli using an oddball paradigm during tonic and phasic periods. An evoked brain potential during REM sleep, the latency of which is approximately 200 ms (hereafter, P2), is elicited to deviant tones (Bastuji et al., 1995a; Sallinen et al., 1996; Takahara et al., 2002; 2006b). Previous studies found that the amplitude of P2 reflected the degree of vigilance during REM sleep. Therefore, we used the P2 amplitude to test whether vigilance was enhanced in one hemisphere with the FNE analogously to NREM sleep. We also tested whether the impact of the FNE on brain responses differed between the phasic and tonic periods because the FNE may differentially influence these periods.

## Materials and Methods

### Participants

Sixteen subjects participated in the present study. Eight subjects participated in Experiment 1 (6 females; 22.3 ± 0.84 yrs, mean ± SEM), and the other eight subjects participated in Experiment 2 (4 females; 24.4 ± 0.86 yrs, mean ± SEM). An experimenter thoroughly described the purpose and procedures of the experiment to candidate subjects, and they were asked to complete questionnaires about their sleep-wake habits for screening, including usual sleep and wake times, regularity of their sleep-wake habits and lifestyle, habits of nap-taking, and information on their physical and psychiatric health, including sleep complaints. The exclusion criteria included a physical or psychiatric disease, currently receiving medical treatment, suspected sleep disorder, and habits of consuming alcoholic beverages before sleep or smoking. Eligible people had regular sleep-wake cycles, i.e., difference between average bedtimes and sleep durations on weekdays and weekends was less than 2 h, and the average sleep duration ranged from 6 to 9 h regularly. All subjects gave written informed consent for their participation in experiments. Data collection was performed at Brown University. The institutional review board approved the research protocol.

Two subjects’ data from day 1 and another 2 subjects’ data from day 2 were excluded from Experiment 2, resulting in the total number of data 6 each for days 1 and 2. The data omitted from day 1 and day 2 were different subjects. The reasons for data omission were lack of REM sleep (n=2), lack of sound presentation due to arousals during REM sleep (n=1), or the measured brain responses were too noisy (n=1).

### Experimental design

Subjects in Experiments 1 and 2 were instructed to maintain their regular sleep-wake habits before the experiments started, i.e., their daily wake/sleep time and sleep duration until the study was over. The sleep-wake habits of the subjects were monitored using a sleep log for 3 days prior to the experiment. Subjects were instructed to refrain from alcohol consumption, unusual excessive physical exercise, and naps on the day before the sleep session. Caffeine consumption was not allowed on the day of experiments.

#### Experiment 1

We tested whether there was hemispheric asymmetry in theta activity in the DMN during REM sleep with the FNE. All the eight subjects took a nap for the first time in the sleep laboratory. Subjects came to the experimental room at approximately 1 pm for PSG preparations. Room lights were turned off at approximately 2 pm, and the 90-min sleep session began. The time for the sleep session was chosen due to the known “mid-afternoon dip”, which should facilitate the onset of sleep even in subjects who do not customarily nap (Horikawa et al., 2013). Structural MRI was measured on another day after the sleep session (see ***Anatomical MRI acquisition and region of interest***).

#### Experiment 2

We measured the evoked brain responses in each hemisphere during REM sleep using an oddball paradigm. We tested whether the evoked brain responses to deviant sounds were larger on day 1 than day 2. Eight subjects participated in two experimental sleep sessions (day 1 and day 2). These sessions were performed approximately one week apart so that any effects of napping during the first sleep session would not carry over to the second sleep session.

Subjects came to the experimental room at approximately 1 pm on both days, and electrodes were attached for PSG measurements (see ***PSG measurement***). Subjects were taken to a sleep chamber after electrode attachment. Subjects were informed that faint beep sounds may be presented through earphones while they slept, but they were instructed to ignore these sounds. Room lights were turned off at approximately 2 pm, and the 90-min sleep session began, as in Experiment 1. PSG was monitored, and experienced experimenters scored sleep stages in real time during the sleep session. Sound presentation (see ***Auditory stimuli***) began after at least 5 min of an uninterrupted NREM sleep stage 2. Sound presentations were stopped every time a lighter stage (stage W or NREM sleep stage 1) was observed. This procedure was repeated throughout the sleep session.

### PSG measurement

PSG was recorded in a soundproof and shielded room. PSG consisted of EEG, electrooculogram (EOG), electromyogram (EMG), and electrocardiogram (ECG). EEG was recorded at 64 scalp sites, according to the 10% electrode position (Sharbrough et al., 1991), using active electrodes (actiCap, Brain Products, LLC) with a standard amplifier (BrainAmp Standard, Brain Products, LLC). The online reference was Fz, and it was re-referenced to the average of the left and right mastoids offline after the recording. The sampling frequency was 500 Hz. The impedance was kept below 20 kΩ because the active electrodes included a new type of integrated impedance converter, which allowed the transmission of the EEG signal with significantly lower levels of noise than traditional passive electrode systems. The data quality with active electrodes was as good as 5 kΩ using passive electrodes, which were used for EOG and EMG (BrainAmp ExG, Brain Products, LLC). Horizontal EOG was recorded from 2 electrodes placed at the outer canthi of both eyes. Vertical EOG was measured from 2 electrodes 3 cm above and below both eyes. EMG was recorded from the mentum (chin). ECG was recorded from 2 electrodes placed at the right clavicle and the left rib bone. The impedance was kept below 10 kΩ for the passive electrodes. Brain Vision Recorder software (Brain Products, LLC) was used for recording. The data were filtered between 0.1 and 40 Hz.

### Sleep-stage scoring and sleep parameters

Sleep stages were scored for every 30-s epoch, following standard criteria (Rechtschaffen and Kales, 1968; Iber et al., 2007), into stage wakefulness (stage W), NREM stage 1 sleep (stage N1), NREM stage 2 sleep (stage N2), NREM stage 3 sleep (stage N3), and REM sleep. The following variables were calculated for each subject for assessment of basic sleep structure and confirmation of the FNE: the duration of each sleep stage, percentage of stage N3 (stage N3%), latency to REM sleep (min), sleep onset latency (SOL, min), wake time after sleep onset (WASO, min), sleep efficiency (SE, %), and time in bed (TIB, min).

### Anatomical MRI acquisition and region of interest

Anatomical MRI data in Experiment 1 were acquired and used to determine the conductor geometry for the boundary element model (BEM) of the head (Hamalainen and Sarvas, 1989) and registering the EEG sensor locations with the individual subject’s anatomy (Dale et al., 1999; Fischl et al., 1999). Subjects were scanned in a 3T MR scanner (Trio, Siemens) using a 32-ch head coil. T1-weighted MR images (MPRAGE; TR = 2.531 s, TE = 3.28 ms, flip angle = 7°, TI = 1100 ms, 256 slices, voxel size = 1.3 × 1.3 × 1.0 mm) were acquired. The cortical surface was inflated for each subject for brain parcellation to localize individual gyri and sulci (Fischl et al., 2004a).

The locations of the DMN were anatomically determined *a priori* based on previously published papers using an automated parcellation method individually (Fischl et al., 2004b; Destrieux et al., 2010; Tamaki et al., 2016). We defined the DMN as a circuit that included the medial prefrontal, inferior parietal, and posterior parietal cortices, according to previous research (Mason et al., 2007; Raichle and Snyder, 2007; Tamaki et al., 2016). The medial frontal cortex consists of the anterior part of the superior frontal gyrus, and the anterior cingulate gyrus and sulcus. The inferior parietal cortex consists of the inferior parietal gyrus and angular gyrus. The posterior parietal cortex consists of the precuneus gyrus, posterior-dorsal cingulate gyrus, and sup-parietal sulcus.

### Source localization of EEG

To compute the strength of brain activities during sleep in the DMN, EEG data were subjected to the Morlet wavelet analysis in Experiment 1 and source localization using the minimum-norm estimate (MNE) of individual MRI information (see ***Anatomical MRI acquisition***). The Morlet wavelet analysis was applied to raw EEG data (Lin et al., 2004; Ahveninen et al., 2007; Tamaki et al., 2013) every 30 s to obtain the MNE strength at the peak frequency of 6 Hz (theta activity) during REM sleep. The window width in the Morlet wavelet analysis was specified as 10, such that the MNE strength would cover the activities from 5-7 Hz, which corresponds to the theta band. To localize the current sources underlying the EEG signals, the cortically constrained MNE was used on EEG using individual anatomical MRI and constrained the current locations to the cortical mantle (Lin et al., 2004; Ahveninen et al., 2007). Information from the EEG sensor locations and the structural MRI segmentation were used to compute the forward solutions for all source locations using a three-layer model of boundary element method (BEM) (Hamalainen and Sarvas, 1989). The individual forward solutions constituted the rows of the gain (lead-field) matrix. The noise covariance matrix was computed from raw EEG data for 30 s during wakefulness. These 2 matrices were used to calculate the inverse operator to yield the estimated source activity during sleep, as a function of time, on a cortical surface (Lin et al., 2004; Ahveninen et al., 2007).

### Classification of phasic and tonic periods

To classify the EEG recordings during REM sleep into phasic or tonic periods in Experiment 2, we first detected eye movements during REM sleep automatically. First, a bandpass filter (0.5-8 Hz) was applied to the vertical and horizontal EOG electrodes during REM sleep. We then measured the amplitudes of eye movements. If the amplitude was 20 µV or greater, then it was counted as 1 movement. Eye movements that occurred within a brief time window (within 100 ms) were counted as only 1 movement. Based on the eye movements, the recordings during REM sleep were classified into a phasic or tonic period every 3 s according to previous studies (Takahara et al., 2002; 2006b). If at least one eye movement was detected within a 3-s epoch, it was classified as a phasic period, and if no eye movement was detected within a 3-s epoch, it was classified as a tonic period.

### Auditory stimuli

Auditory stimuli were controlled using MATLAB (The MathWorks, Inc.) software and presented through earphones (HAFR6A, JVC Americas Corp.). The stimuli consisted of 2000-Hz deviant (presented at 10% probability) and 1000-Hz standard (presented at 90% probability) pure tones, all 50 ms in duration (10 ms rise/fall). These sounds were presented monaurally every 1 s (1 trial = 1000 ms, fixed ISI = 950 ms). The probabilities of sound type (deviant or standard) and the presented ear (left or right) were pseudorandomized every 30 s, which corresponded to the sleep-stage scoring epoch (see ***Sleep-stage scoring***). More concretely, 30 sounds were presented in total, 15 per ear, in a given 30-s epoch. The probability of a deviant sound was 10%, and at least one deviant sound was presented to each ear.

The sound intensity was approximately 35 dB (Extech 407740, Digital Sound Level Meter, Extech Instruments Corp.), which was lower than previous studies that also used an oddball paradigm during REM sleep (50-100 dB) (Sallinen et al., 1996; Cote and Campbell, 1999; Cote et al., 1999; Cote et al., 2001; Takahara et al., 2002; 2006b) to prevent subjects from waking. It was confirmed that 35 dB was sufficiently quiet to maintain sleep for each subject before the sleep session began.

### Analysis of brain responses to auditory stimuli

We examined the evoked brain potential known as the P2 (Takahara et al., 2002; 2006b) from EEG data recorded during REM sleep using an oddball paradigm in Experiment 2. P2 is a positive brain potential that appears during REM sleep, and its amplitudes to rare and salient stimuli increase (Takahara et al., 2002; 2006b). Therefore, these responses are used as an index of vigilance during REM sleep in humans.

To obtain the P2, 6 frontal channels (3 channels per hemisphere) were analyzed (left: FC1, FC3, FC5; right: FC2, FC4, FC6). The frontal region was chosen for the current analysis because these areas are near the DMN, where interhemispheric asymmetry was observed with the FNE during deep NREM sleep (Tamaki et al., 2016). All data were examined visually for each trial, and any trials that included arousals (Bonnet, 1992; Iber et al., 2007) or motion artifacts were excluded from further analyses.

Analyses of brain responses followed a previous study (Tamaki et al., 2016). The amplitudes of EEG during the 200-ms prestimulus period (−200 ms to 0 ms) were averaged. The mean EEG amplitude of the prestimulus period was subtracted from the EEG amplitudes from the 0- to 1000-ms poststimulus period to normalize the signal amplitude for each of the 1-s trials (0 to 1000-ms poststimulus). These normalized values were averaged for each sound type (deviant and standard), hemisphere (left and right), day (day 1 and day 2), and period (tonic and phasic) during REM sleep to compute averaged brain responses. The maximum value (peak) of the 150-250-ms poststimulus time window from the brain responses was used as the P2 amplitude. We chose this time window because it roughly corresponds to the defined P2 in previous studies (Bastuji et al., 1995b; Perrin et al., 1999; Crowley and Colrain, 2004; Takahara et al., 2006b). Averaged values were obtained for each of the phasic and tonic periods (see ***Classification of phasic and tonic periods***).

### Statistical analyses

The α level (type I error rate) of 0.05 was set for all statistical analyses. In Experiment 1, a two-tailed paired t-test was performed for analyses of theta activity. In Experiment 2, a 3-way repeated measures ANOVA was used for the analysis of P2. In post hoc tests, t-tests were used with Bonferroni correction.

## Results

### Confirmation of the FNE

To confirm that the FNE occurred in Experiments 1 and 2, we compared the sleep-onset latency, WASO, and stage N3% on day 1 in Experiments 1 and 2 to day 2 in Experiment 2. We confirmed that the FNE occurred in both experiments (**Table 1**). We found that the sleep-onset latency, which is a critical measure for the FNE (Schmidt and Kaelbling, 1971; Webb and Campbell, 1979b; Tamaki et al., 2005a), was significantly longer on day 1 than day 2 (Exp. 1: t (14) = 3.25, p = 0.006; Exp. 2: t (7) = 3.60, p = 0.009). We also found that WASO was significantly larger (Exp. 1: t (14) = 2.17, p = 0.048; Exp. 2: t (7) = 3.21, p = 0.015) and stage N3% was significantly lower (Exp. 1: t (14) = 2.57, p = 0.022; Exp. 2: t (7) = 2.41, p = 0.047) on day 1 than day 2. These results indicate that it took a longer time to fall asleep on day 1 compared to day 2, sleep was more fragmented, and deep sleep was reduced, in accordance with previous studies (Kales et al., 1967a; Scharf et al., 1975; Webb and Campbell, 1979a; Tamaki et al., 2016). However, there was no significant difference in the duration of (Exp. 1: t (14) = 0.76, p = 0.461; Exp. 2: t (7) = 1.03, p = 0.337) or latency to (Exp. 1: t (13) = 0.75, p = 0.466; Exp. 2: t (7) = 0.43, p = 0.676) REM sleep. See **Table 1** for the results of all the sleep parameters for Experiments 1 and 2.

**Table 1.**
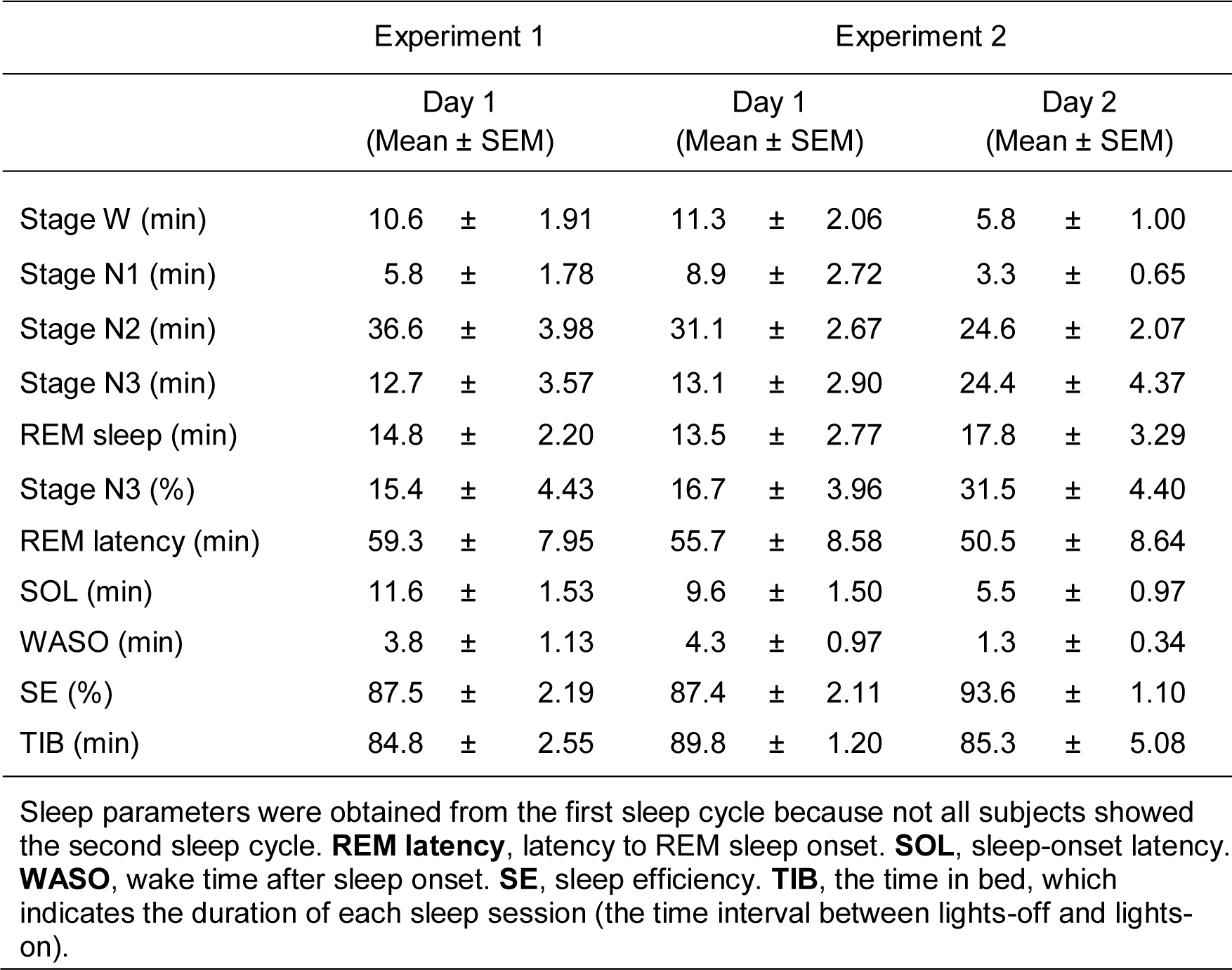
Sleep parameters

#### Experiment 1

##### Source-localized theta activity

Experiment 1 examined whether there was interhemispheric asymmetry in theta activity in the DMN during REM sleep in association with the FNE. We measured theta activity originating in the DMN in each hemisphere (see ***Source localization of EEG*** in **Methods**). We performed a two-tailed paired t-test on measured theta strength between the left vs. right hemispheres. The result of the t-test was not significant (t (7) = 0.1, p = 0.938; **Figure 1**).

**Figure 1.**
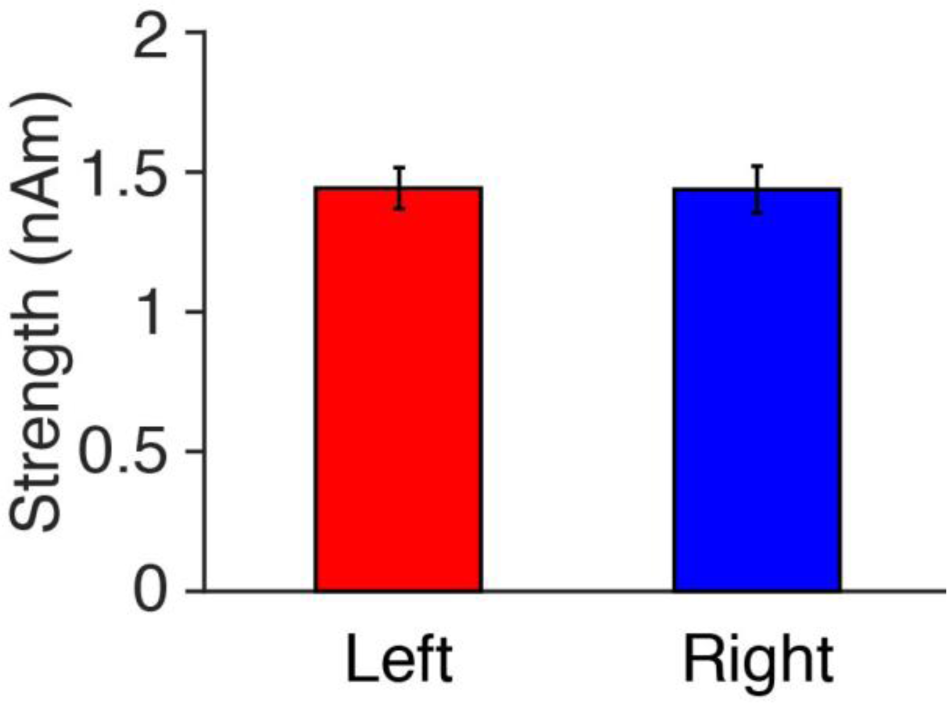
Source-localized theta strength in DMN in the left and right hemispheres. N=8 in total.

#### Experiment 2

##### Evoked brain response

We obtained the P2 brain potential (**Figure 2**), which is one of the primary evoked components during REM sleep (Takahara et al., 2002; 2006b). The amplitude of P2 may differ by eye movement state. Therefore, we analyzed the brain responses for the tonic (no REMs) and phasic (1 or more REMs) periods (see ***Classification of phasic and tonic periods*** in **Methods**).

**Figure 2.**
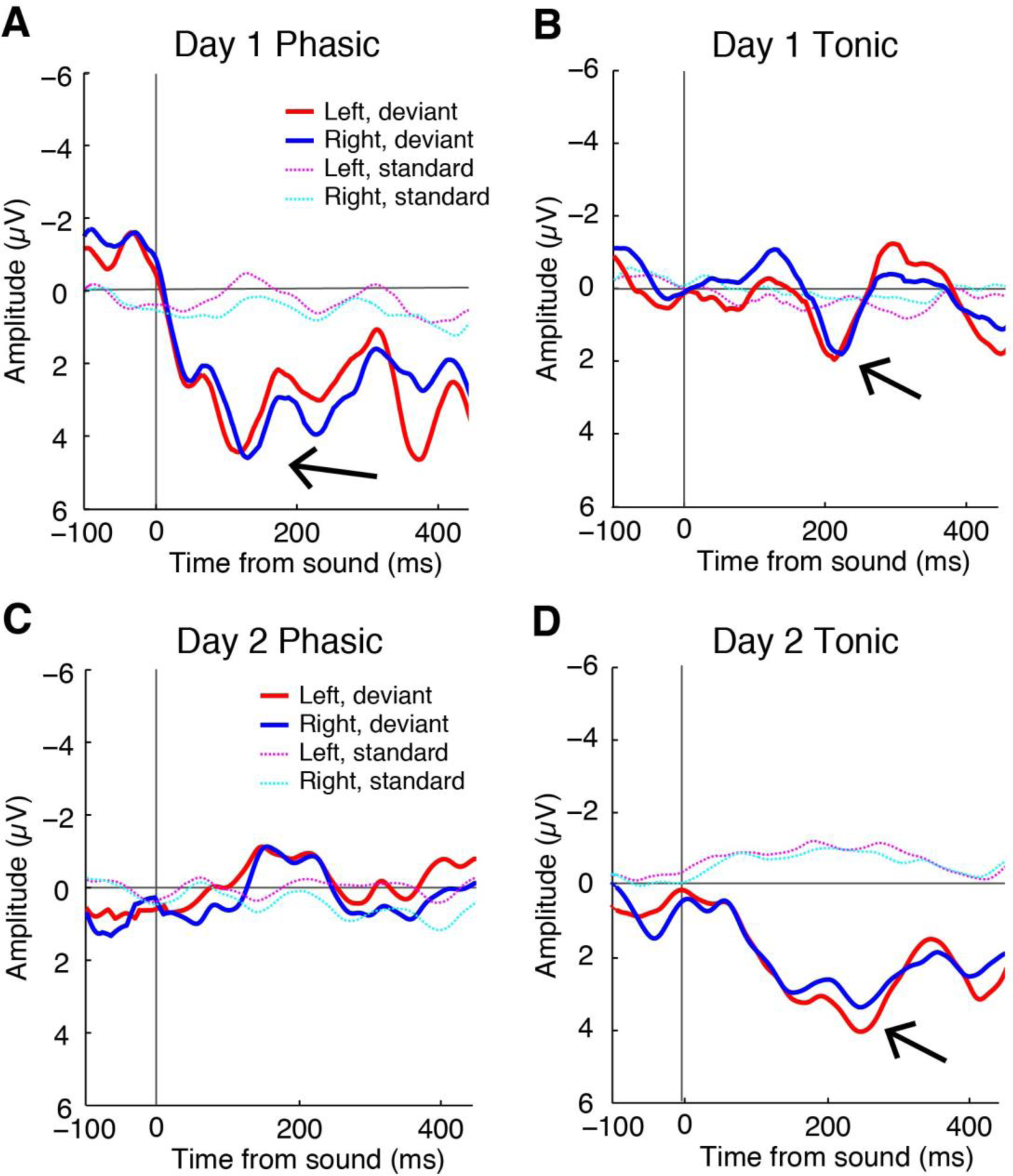
The grand-averaged brain responses to deviant sounds time-locked to the sound onset during the phasic (**A**) and tonic (**B**) periods on day 1 (N=6), and the phasic (**C**) and tonic (**D**) periods on day 2 (N=6) during REM sleep. The data in the figures are smoothed for visualization purposes via application of a moving average (20 data points window). Arrows indicate P2.

To test whether the brain was vigilant in a brain hemisphere during REM sleep with the FNE, and whether the vigilance level differed by eye movement state, we measured the mean amplitudes of the evoked brain responses for each hemisphere (left vs. right), period (phasic vs. tonic), and sound (deviant vs. standard) (see ***Analysis of brain responses to auditory stimuli*** in **Methods** for measurement of amplitude). We found that P2 was elicited only during the tonic period, and not during the phasic period, on day 2 (**Figures 2C and D** for the ground-averaged brain responses on day 2, which replicates previous studies). However, a large P2 was elicited to deviant tones during tonic and phasic periods on day 1 with the FNE (**Figures 2A and B** for the ground-averaged brain responses on day 1). However, P2 was elicited only during the tonic period on day 2, and not during the phasic period (**Figures 2C and D** for the ground-averaged brain responses on day 2).

A 3-way ANOVA with within-subjects factors of Period (tonic, phasic) and Hemisphere (left, right) and a between-subjects factor of Day (day 1, day 2) was performed on the P2 amplitudes elicited by deviant tones (**Figure 3**). The Day factor was a between-subject factor due to data omission (see ***Participants*** above). If there was interhemispheric asymmetry in the brain response associated with the FNE, then a significant Day x Hemisphere interaction should occur. If the amplitudes were different between days in association with the period, then a significant Day x Period interaction should occur.

**Figure 3.**
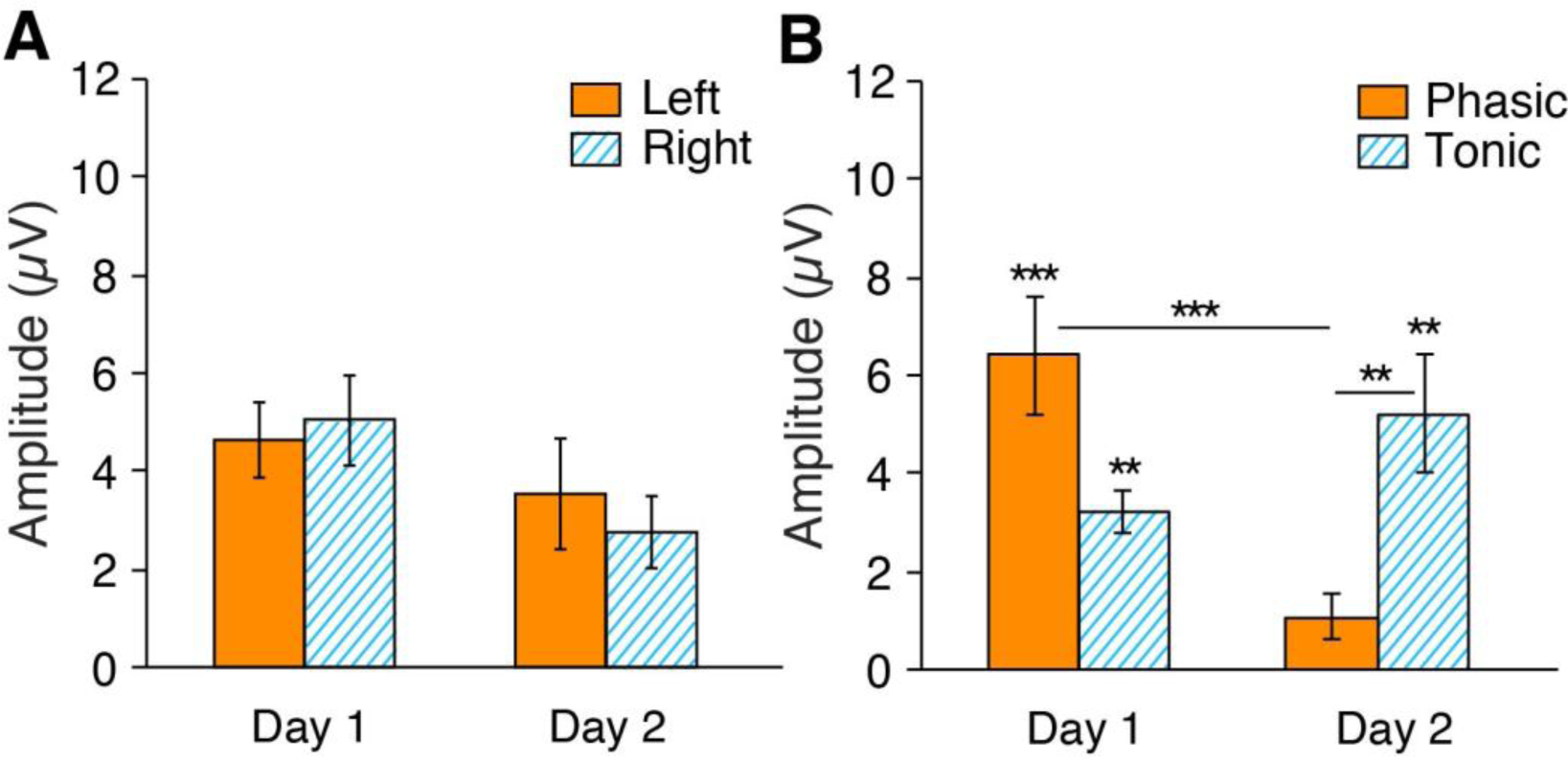
The mean P2 amplitudes to deviant sounds during REM sleep. (**A**) The amplitudes for each hemisphere on days 1 and 2 (phasic and tonic periods averaged). (**B**) The amplitudes for each phasic and tonic period on days 1 and 2 (hemispheres averaged). N=6 for day 1 and n=6 for day 2. Post hoc t-test, ***p<0.001, **p<0.01 (Bonferroni correction).

The statistical results showed that the Day x Hemisphere interaction was not significant (**Figure 3A**; F (1, 20) = 0.56, p=0.465). However, there was a significant Period x Day interaction (**Figure 3B**; F (1, 20) = 11.56, p = 0.003). None of the main factors (Hemisphere, F (1, 20) = 0.07, p=0.791; Period, F (1, 20) = 0.19, p=0.668; Day, F (1, 20) = 2.41, p=0.136) or other interactions (Hemisphere x Period, F (1, 20) = 1.28, p=0.272; Day x Hemisphere x Period, F (1, 20) = 0.06, p=0.810) were significant. Because a Period x Day interaction was significant, we performed post hoc analyses to investigate the source of this interaction. Because no significant effect of Hemisphere was found in the above ANOVA, data from the left and the right hemispheres were pooled for subsequent analyses. Post hoc t-tests indicated a significant difference in the amplitude between day 1 vs. day 2 in the phasic period (**Figure 3B**; unpaired t-test, t (24) = 4.31, p = 0.002, Bonferroni correction for the 4 comparisons) and between the phasic and tonic periods on day 2 (**Figure 3B**; paired t-test, t (11) = 3.30, p = 0.029). There was no significant difference between days in the tonic period (**Figure 3B**; unpaired t-test, t (22) = 1.57, p = 0.131) or between the tonic vs. phasic periods on day 1 (**Figure 3B**; paired t-test, t (11) = 2.29, p = 0.171). We performed one-sample t-tests on the amplitudes for each of the periods and days to investigate whether P2 was elicited, being larger than zero, on days 1 and 2 for the phasic and tonic periods. (**Figure 3B**). The amplitude was significantly different from 0 during the tonic periods on day 1 (t (11) = 7.92, p < 0.001, Bonferroni correction for the following 4 comparisons) and day 2 (t (11) = 4.34, p = 0.005), and during the phasic period on day 1 (t (11) = 5.29, p < 0.001), but not on day 2 (t (11) = 2.30, p = 0.168).

## Discussion

The present study found that the vigilance level was higher on day 1 than day 2, specifically during the phasic period. However, interhemispheric asymmetry in evoked brain responses was not found during REM sleep. Theta activity during REM sleep did not show interhemispheric asymmetry.

Notably, although the brain responses to auditory stimuli did not show interhemispheric asymmetry associated with the FNE, a larger response to rare stimuli was found, specifically during the phasic period. Because the amplitude in an evoked brain response to deviant stimuli correlates with the degree of vigilance (Nielsen-Bohlman et al., 1991; Michida et al., 2005), the present results suggest that the FNE augments vigilance during REM sleep, especially when rapid eye movements are observed. These results demonstrate that a night-watch system using both hemispheres exists during the phasic period of REM sleep.

We found that the amplitudes of brain responses were larger on day 1 than day 2, specifically during the phasic period. Why did the phasic period, but not the tonic period, show augmented evoked brain responses in association with the FNE? The brain is more sensitive in the monitoring of external stimuli during the tonic period than the phasic period during normal sleep without the FNE (Takahara et al., 2002). Therefore, the capacity for information processing or attention to external stimuli may already be limited (Kahneman, 1973; Marois and Ivanoff, 2005) during the tonic period. If the attentional resource is spared for external monitoring during the tonic period, then when there is a need to increase vigilance even further during sleeping in an unfamiliar environment, such an increase would have to occur outside the tonic period, i.e., during the phasic period.

The phasic period during normal REM sleep without the FNE is linked to subjective mental activities (Berger and Oswald, 1962; Weinstein et al., 1988). Notably, a previous study suggested that the sensitivity to external stimuli was lowered when there is sleep-onset dreaming (Michida et al., 2005). Therefore, the reason that external monitoring is impaired during the phasic period of normal REM sleep without the FNE may be due to ongoing mental activities, including dreaming, which may interfere with the monitoring of the external environments (Sallinen et al., 1996; Michida et al., 2005). However, the large brain response was elicited during the phasic period during REM sleep in association with the FNE. This result suggests that resources for internal mental activities are deployed for the external monitoring during sleeping in an unfamiliar environment when the FNE occurs.

We did not find a clear interhemispheric asymmetry in theta activity or the amplitude of the evoked potentials between hemispheres during REM sleep in association with the FNE. These results contrast our previous study (Tamaki et al., 2016) in which we found interhemispheric asymmetry in regional slow-wave activity and vigilance during deep NREM sleep. This difference suggests that a different mechanism than deep NREM sleep applies to a night-watch system during REM sleep. The present results showed that the vigilance to deviant sounds increased on day 1, which was associated with the FNE. Therefore, a type of surveillance to the external world may exist during REM sleep. It may be the case that the arousal threshold may be too high and costly during deep NREM sleep to increase vigilance in both hemispheres, and only one hemisphere may be used for surveillance. However, there are already resources for dreaming and mental activity during REM sleep. These resources may be used for surveillance in both hemispheres. This deployment of resources may occur without much sacrifice in sleep depth during REM sleep.

In conclusion, REM sleep has a protective mechanism that may involve a different mechanism than deep NREM sleep. A night-watch during REM sleep was shown as increased vigilance in both hemispheres throughout REM sleep, specifically during the phasic period. This REM-specific night-watch system may be realized by deploying the resources available to internal activity for surveillance during sleeping in an unfamiliar environment.

## Acknowledgements

This work was supported by NIH (R21EY028329) and Institutional Development Award (IDeA) from the National Institute of General Medical Sciences of the National Institutes of Health (P20GM103645), Center for Vision Research Brown University, and FY17 OVPR Seed Grant, Brown University.

## Author Contributions

M.T. and Y.S. designed the research and wrote the manuscript. M.T. performed the experiments and analyzed the data.

